# Neuromechanical phase lag predicts material and neural control properties in *Caenorhabditis elegans*

**DOI:** 10.1101/312389

**Authors:** Jack E. Denham, Thomas Ranner, Netta Cohen

**Affiliations:** School of Computing, University of Leeds, UK

**Keywords:** pattern generation, proprioception, motor control, biomechanics, undulatory locomotion, nematodes

## Abstract

In all animals, the successful orchestration of motor programs hinges on appropriate coupling between components of the system, from neural circuit dynamics, through muscles and body properties to the physical environment. We study this coupling in undulatory locomotion, with a view to better understanding the relative roles of central and reflex-driven control. We ask how the coupling between neural control and body mechanics is affected by sensory inputs during undulatory locomotion in *C. elegans.* To address this question, we use a biomechanical simulation framework, within which we separately model feed forward and feedback controlled undulations. We characterize neuromechanical phase lag and locomotion speed using body stiffness as a control parameter. We show that sensory entrainment can suppress neuromechanical phase lag, that would otherwise emerge under centrally generated feed forward control.

## INTRODUCTION

Understanding neural circuits that allow animals to orchestrate and fine tune their locomotion behaviors is a long standing endeavor. A variety of behaviors studied to date, from human gait to insect stepping, and from lamprey swimming to locust flight, tend to be controlled by distributed central pattern generating (CPG) circuits that are subject to activation and modulation by descending control and ascending sensory information,^12,18^ often in the form of proprioception or information about parts of the body, their position and movement. The roles of proprioception vary greatly across different systems.^4,13,14,19,24^ Indeed in a number of cases, sensory neurons are embedded within the pattern generating circuit itself, such that a meaningful description of the behaviorally relevant pattern generation mechanism invariably combines the two.^1^ Whatever the rhythm generating mechanism, however, all motor behavior is ultimately generated by the combined action of neural circuits and a physical system that includes muscles embedded in a body that interacts with an environment. It stands to reason, therefore, that an integrated understanding of neural circuits and biomechanics can provide a much more complete understanding of the control of motor behavior.^5,21^

One manifestation of the coupling between neural circuits and the body which is evident in a variety of swimming animals is an advancing neuromechanical phase.^8,15,22^ When a retrograde undulation is generated, a neuromech-nical wave propagates from head to tail, resulting in forward thrust of the body. Often, the wave speed of neuromuscular activation is faster than that of body mechanics, resulting in an advancing neuromechanical phase lag: a phase lag between the neural and muscle activation on the one hand and the mechanical action of the body on the other, that grows from head to tail. The wide range of animal sizes and conditions in which neuromechanical phase lags arise suggests that such phase lags are fundamental to undulatory locomotion.

Much of our insight into neuromechanical phase lag has come from mathematical and robotic models. In a model of lamprey, Tytell *et al.^22^* present a thorough analysis of phase lags, the conditions for their emergence, the accompanying body shapes and the consequences for locomotion, including speed, the power needed and capacity for rapid acceleration. Here, as in a variety of fish, neuromechanical phase lag arises when the forces produced by the body wall muscles are weaker than those applied by the surrounding water. A model of sandfish, desert lizards that ‘swim’ through granular fluids,^8^ reports an entrainment of bending torque between distant parts of the body. In all these studies, the models focused on feed forward, open loop control of the bending of the body. Thus the contribution of sensory, proprioceptive feedback to shaping neuromechanical phase lags remains poorly understood. Here, we revisit the question of neuromechanical phase lag by comparing feed forward and feedback (closed loop) control within a neuromechanical simulation framework of *Caenorhabditis elegans.*

*C. elegans* is a leading neurobiological model system, with a large body of experimental and modeling locomotion work, and yet the question of neuromechanical phase lag has never before been addressed in this system. This nematode worm undulates by alternating dorsal and ventral contractions that propagate rhythmically from head to tail. Muscle contractions are controlled by motor neurons that line the ventral nerve cord. *C. elegans* moves effectively in a wide range of external fluid and complex heterogeneous environments. When placed in different environments, *C. elegans* aptly adjusts its waveform and undulation frequency.^2,9^ Such gait modulation is characterized by a gradual decrease in both the frequency and wavelength of undulations with growing environmental drag forces, with the likely benefit of optimizing locomotion speed.^6^ Experimental and computational models clearly point to the importance of incorporating active muscles, mechanical coupling and the environment to recapitulate the kinematic parameters as well as the observed gait modulation.^3,20^

While the neural circuitry of *C. elegans* is fully mapped, the pattern generating mechanism responsible for locomotion remains mysterious. On the one hand, the evidence to date suggests a number of complementary proprioceptive pathways and functions. Previous computational models as well as experimental studies suggest that within the ventral nerve cord, proprioception may be sufficient for pattern generation and is necessary for coordinating dorso-ventral anti-phase contractions as well as for imposing the appropriate wave speed along the body.^3,11,23^ Recent papers also present support for CPGs within this circuit^10,11^ but the detailed mechanisms for either endogenous oscillations or proprioception are still to be determined.

Here, we use a modeling approach to compare CPG and reflex driven control. We focus on two metrics of the locomotion: the neuromechanical phase lag, and the locomotion speed. In doing so, we seek experimental predictions that may shed light on the dominating mechanism for pattern generation in *C. elegans*. As in the lamprey model^22^ we find that advancing neuromechanical phase lags arise naturally from centrally generated feed-forward control in biomechanical systems characterized by a relatively low effective bulk Young’s modulus. In contrast, in a model where proprioception dominates the pattern generation, we find that body shape entrains the neural activation, thus considerably suppressing phase lags along the body. We compare the kinematics of locomotion associated with these systems for different profiles of neuromechanical phase lag, pointing out subtle behavioral differences that may serve as signatures of different mechanisms of neural control. Finally, we consider the locomotion speed under the variety of conditions tested and show that the form of control has a drastic effect on the locomotion speed across a range of conditions and body properties.

## MODELS

### Mechanical model

Here we use a continuum mechanical model subject to resistive force theory to capture the drag forces describing the worm’s low Reynolds number undulatory movement^6^ (Fig. 1a). The worm’s body is represented by a thin viscoelastic shell. The high internal pressure in *C. elegans* is represented as a line tension *p* along this midline and is chosen such that the midline is inextensible (and set to 1 mm from head to tail). Internal pressure helps maintain the worm’s shape and relaxes the body back to a straight configuration in the absence of muscle activation. The *C. elegans* cuticle is innervated by 95 body wall muscles arranged in four quadrants along the body. In the model, bending due to active muscle force is represented by a torque acting on the midline of the body. As we show below, this torque may be expressed as a *preferred curvature β* = *β(u,t)* along the body midline; here, *u* denotes the position along the midline of the body (from 0 to 1) and t denotes time. Thus, *β* will vary along the worm and in time, with positive and negative values corresponding to ventral or dorsal excitation respectively. The body curvature, *κ* = *κ*(*u,t*), generated by the active moment, follows the muscle forcing, *β*(*u,t*), on a timescale proportional to muscle strength and Young’s modulus *E*, which acts much like a spring constant to resist displacement. Finally, the resistive environmental drag forces

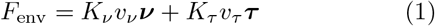

are decomposed into two forces acting in directions normal and tangential to the body, with corresponding drag coefficients *K_ν_ ≥ K_τ_* acting along the directions *ν* and *τ* respectively; here, *υ_ν_* and *υ_τ_* denote the normal and tangential components of the speed of the body. The balance of forces is summarized as follows:

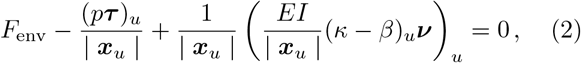

where *I* is a geometrical entity, *x* denotes a coordinate of a point along the body (in the lab frame) and the subscript *u* denotes the derivative with respect to *u* (along the midline of the worm). Equation (2) conveniently allows us to seamlessly translate between units of torque and body curvature. Thus, we may represent the torque as a preferred curvature *β(u,t)*. Zero force and zero torque are enforced at the boundaries, such that *β = κ* at both ends of the body. The model is solved using a finite element method^6^.

**FIG. 1:**
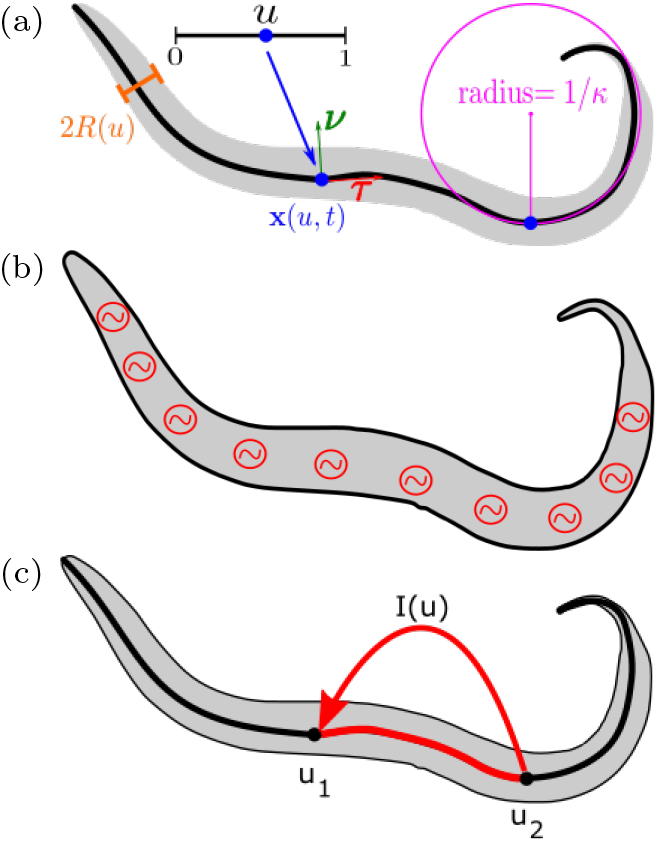
Model schematics: (a) body mechanics (b) feed forward control (c) feedback proprioceptive control.

### Neuromechanical coupling

The neuromuscular component of the model will be split into two cases. First, in the feed forward case, it is assumed that central pattern generators located along the ventral nerve cord are solely responsible for the generation and propagation of an excitation wave (Fig. 1b). Secondly, in the case of feedback driven control, proprioception drives bistable neurons (Fig. 1c). This latter case subsumes to some extent some scenarios in which central pattern generated oscillations are fully entrained to a proprioceptive input.

Both cases include a caricature of the body wall muscles, which are assumed to be continuous along the ventral and dorsal sides, producing torque linearly in response to neural activation. The corresponding linear torque equation is then given by

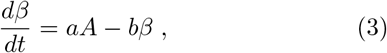

where *A* = *A*(*u,t*) represents a spatiotemporal neural activation, and *β* the torque experienced by the midline as a result of alternating muscle contractions. Constant parameters *a* and *b* respectively determine the sensitivity of muscles to neuromuscular stimulation and the muscle time constant of 1/*b* = 100 ms.

### Central pattern generated control

Centrally generated or feed forward control is modeled by a rhythmic pattern that acts in antiphase on the two sides of the body and travels down the body with an appropriate wave speed. In *C. elegans*, the main motor neurons implicated in forward locomotion are B-type excitatory and D-type inhibitory motor neurons. Both ventral and dorsal motor neurons are disributed along the body (with cell bodies in the ventral nerve cord). Ventral and dorsal motor neurons (of classes B and D) innervate ventral and dorsal body wall muscles, respectively. However, the dynamics of the motor circuit, or of its motor neurons, is not known. Here, we consider a minimal model in which the action of this circuit on the muscles can be captured by continuous unit amplitude oscillations with an imposed period, T, and body wavelength, *λ*, propagating from head to tail. Thus, the input to the muscles is given by traveling sine wave

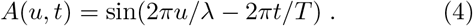

We note that the model is agnostic to the relative contributions of B- and D-type neurons, assuming only that their action on the muscles is phase-synchronized.

In many ways, this toy model is unrealistic. For example the amplitude of undulations typically decays from head to tail in *C. elegans.* Our choice of this model is motivated precisely by its simplicity, to serve as a basis for future studies with more realistic models of neural control.

### Proprioceptively driven control

We assume proprioceptively generated control relies exclusively on feedback from the body shape of the worm. Boyle *et al.^3^* modeled B-type neurons as bistable elements, inspired by electrophysiological recordings of RMD head motor neurons.^16^ In the same model, Boyle *et al.* assumed that ventral D-type neurons are synchronized with dorsal B-type motor neurons and vice versa. Under this assumption, the action of B- and D-class neurons on the same side of the body can be reduced to a single input current function to the muscle, and can therefore be treated as a single bistable element. Here, we similarly treat the neural activation by bistable binary switches. In our continuous representation of the body, the state of ventral and dorsal neurons at a point *u* along the body is given by *V^V^(u), V^D^(u)* ∈ {0,1} with ON (*V* = 1) and OFF (*V* = 0) states.

The input to the body consists of two components: a tonic neural input from the circuit (e.g., from command interneurons) that can be absorbed into a switching threshold *θ*, and a dynamic proprioceptive input current, which we denote *I(u)*. We consider the proprioceptive input as the integral of the body stretch over a section of the body. Due to the fixed width of the body, the contraction of one side of the body results in an equal stretch on the other side, and can be represented compactly by the curvature of the midline. We therefore define the proprioceptive input current 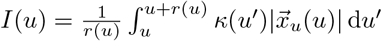 as the integral of midline curvature over a specified proprioceptive range *r(u)*. Here, we adopt a convention whereby a positive input value corresponds to ventral bending, and a negative value to dorsal bending. When the proprioceptive input *I(u)* exceeds some threshold *θ* (a constant parameter along the worm), the dorsal neuron will switch ON, and the ventral one OFF; when the input falls below some threshold (here, taken as –*θ* for symmetry) the dorsal neuron switches OFF and the ventral one ON. Neuronal states are initialized according to the initial body shape with *V^V^(u)* = 0, *V^D^(u)* = 1 if *I*(*u*) > 0 and *V^V^(u)* = 1, *V^D^(u)* = 0 if *I(u)* ≤ 0. Neuronal state switching is then given by

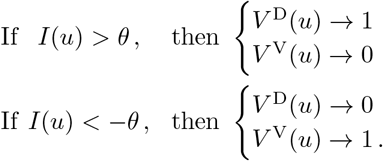

Here, the proprioceptive range is set to half the body length or 0.5 in the anterior half of the body and to 1 – *u* in the posterior half. The activation thresholds are set to an effective curvature of *θ* = 3 mm^−1^. The sensitivity of the system to these parameters is explored in the accompanying paper.^7^ By imposing anti-phase activation, the state at position *u* along the body reduces to the difference between dorsal and ventral activations *A*(*u*) = *V^D^* (*u*) – *V^V^* (*u*) = ±1. This activation *A*(*u*) is directly used as the muscle input in Eq. (3).

## RESULTS

Whether neuromechanical phase lag plays a role in *C. elegans* locomotion depends on a large number of factors, from the material properties of the body to the mechanical load imposed by the fluid environment. Therefore, in all our simulations, we used the Young’s modulus of the body as a control parameter and considered two environmental conditions: swimming in water and crawling through highly viscoelastic environments, mimicking agar.

### Advancing neuromechanical phase lags emerge under feed forward control

We first simulated agar-like conditions. Under feed forward, CPG like control, we set the period of undulations to *T* = 2 sec and the wavelength *λ* = 0.5 mm. For this undulation period, phase lag, *φ*, and time lag, *Δ*, are related by *φ* = 2*πΔ/T* = *π*Δ. Simulations of feed forward control showed that the neuromechanical phase lag increases linearly with body coordinate, but the rate of increase depends strongly on the body stiffness. For Young’s moduli of 0.4 MPa or higher, the phase lag is effectively negligible. For intermediate body stiffness around 70-100 KPa, the advancing phase is clearly significant, with the phase lag growing to approximately a third to a half of an undulation period towards the tail; hence, the corresponding time lags should be directly measurable. At lower values of the Young’s modulus, the phase lag rapidly grows and below 20 KPa, it exceeds a full period of undulation and the worm becomes too flaccid to generate a coordinated undulatory gait. Since neuromechanical phase lags are known to grow with the mechanical load on the animal, we asked to what extent our results are affected by the external medium. Before running the simulation experiments in water, we adjusted the period and wavelength of undulations to *T* = 0.5 sec and *λ* = 1.5 mm to match behavioral observations. We found that the neuromechanical phase lag in water was nearly abolished. The phase lag reached a maximum of approximately 0.1 radians with *E* ≈ 20 KPa, corresponding to a maximal time lag of approximately 10 msec, i.e., much shorter even than the muscle time scale in our model (not shown). Higher body stiffness reduced this lag even further. Thus, in water, even a body with *E* of 20 KPa responds almost instantly to the neuromuscular drive. In all our simulations, the phase lag no longer advanced from head to tail, but peaked mid-way along the body, suggesting that boundary effects dominate in this regime.

### Proprioceptive feedback can clamp neuromechanical phase lags

To find out if feed forward and feedback control yield different magnitudes and spatial profiles of neuromechanical phase lag, we ran simulations for the same values of Young’s modulus subject to feedback control. In contrast to our results above, simulations of proprioceptively driven locomotion on agar nearly abolished the lag entirely in the anterior half of the body, for all values of the body stiffness. Interestingly, for all values of the Young’s modulus, a neuromechanical phase lag also appeared and advanced rapidly in the posterior part of the body. Thus, even for a Young’s modulus as high as 1 MPa, a phase lag of about 1/3 of a period is obtained at the very tail. As in the case of centrally generated, feed forward control, the lower the elasticity, the larger the phase lag observed, but the effect of body stiffness only manifests in the posterior part of the body. We note that unlike our simulations of feed forward control, here the frequency of undulations is an emergent property of the system and its interactions with the environment. Therefore a significant phase lag does not necessarily imply a significant (measurable) time lag. In our simulations, higher body stiffness leads to faster undulations^7^ and hence the smaller the time lag. It is striking that the point along the body at which the phase lag emerges under feedback control appears to coincides with the range of the proprioceptive field (here, half the body length). Since the proprioceptive input is integrated posteriorly to the local muscle, the receptive field is reduced from midway along the worm (*u* ≥ 0.5).^17^ To validate whether this is the case, we re-ran simulations with different values of receptive field and found that indeed, the phase lag emerges consistently at the point along the worm at which the receptive field begins to decrease (not shown).

To interpret our results, we asked how the phase lags related to the propagation of bends along the animal. Figure 3 demonstrates the advancing phase lags, *φ_κ_(u,t)* – *φβ* (*u, t*), for feed forward and feedback control simulation with a Young’s modulus of 100 KPa in agar-like conditions. Under feed forward control, zero contours of both *β* and *κ* show a characteristic and consistent wave speed in both the neuromuscular activation and curvature along the body. In this case, the phase lag emerges because the curvature wave speed is slower than that of the electrical activation. In contrast, under feedback control, the wave speed and wavelength vary along the body. In particular, zero torque (or preferred curvature) contours show a drastic increase in both wavelength and wave speed midway along the body but the curvature follows approximately at a fixed wave speed. In the anterior of the body, the curvature follows the activation closely, with minimal phase lag, whereas in the posterior, the body fails to catch up with the fast neural activation, resulting in a significant, rapidly advancing phase lag.

**FIG. 2:**
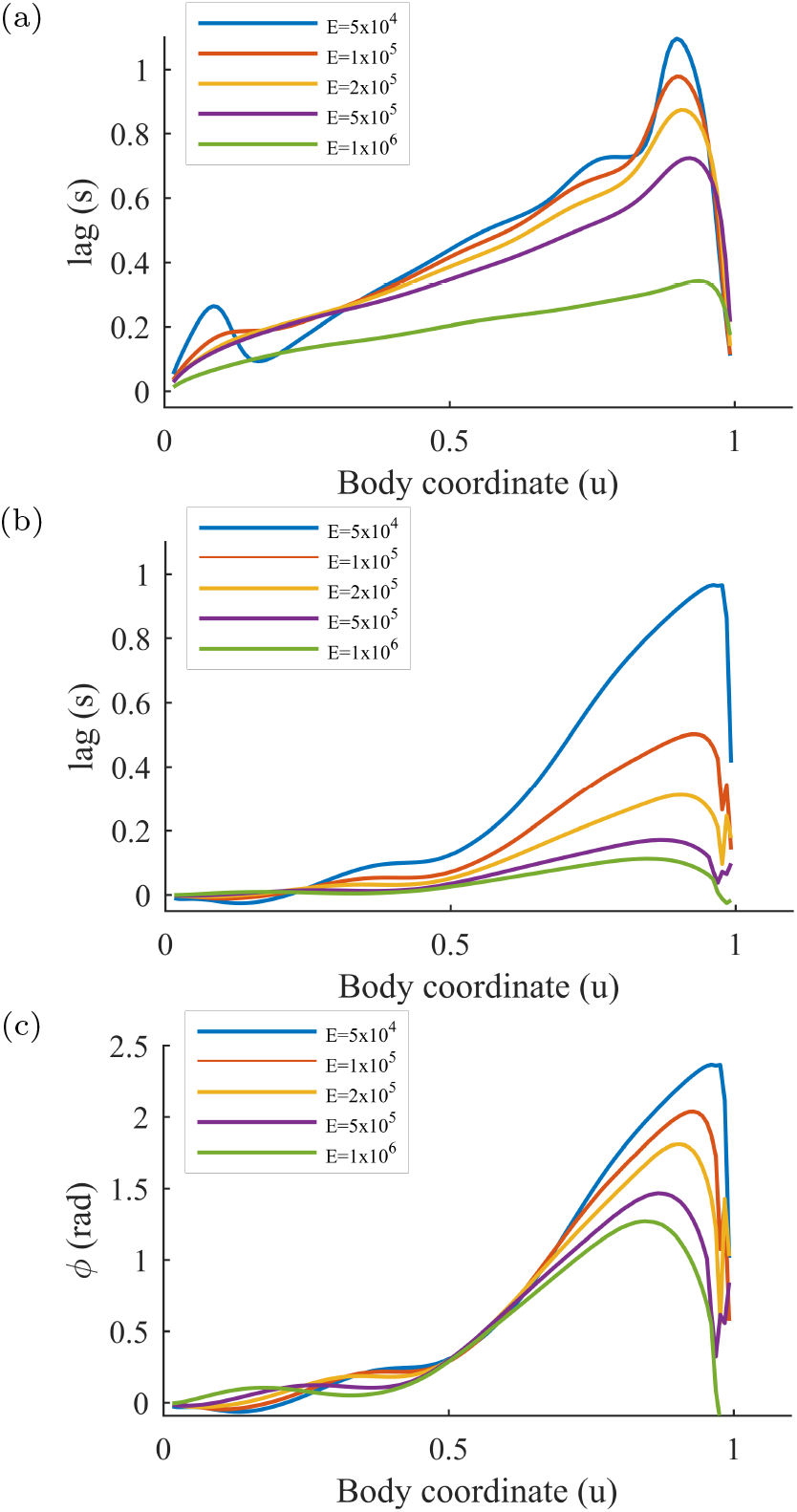
Neuromechanical time lag and phase lag *φ* along the body, for different values of Young’s modulus of the body *E*. (a) Time lags under feed forward control in agar (undulation period 2 sec). (b) Time lags under proprioceptive control in agar. (c) Corresponding phase lags under proprioceptive control in agar.

**FIG. 3:**
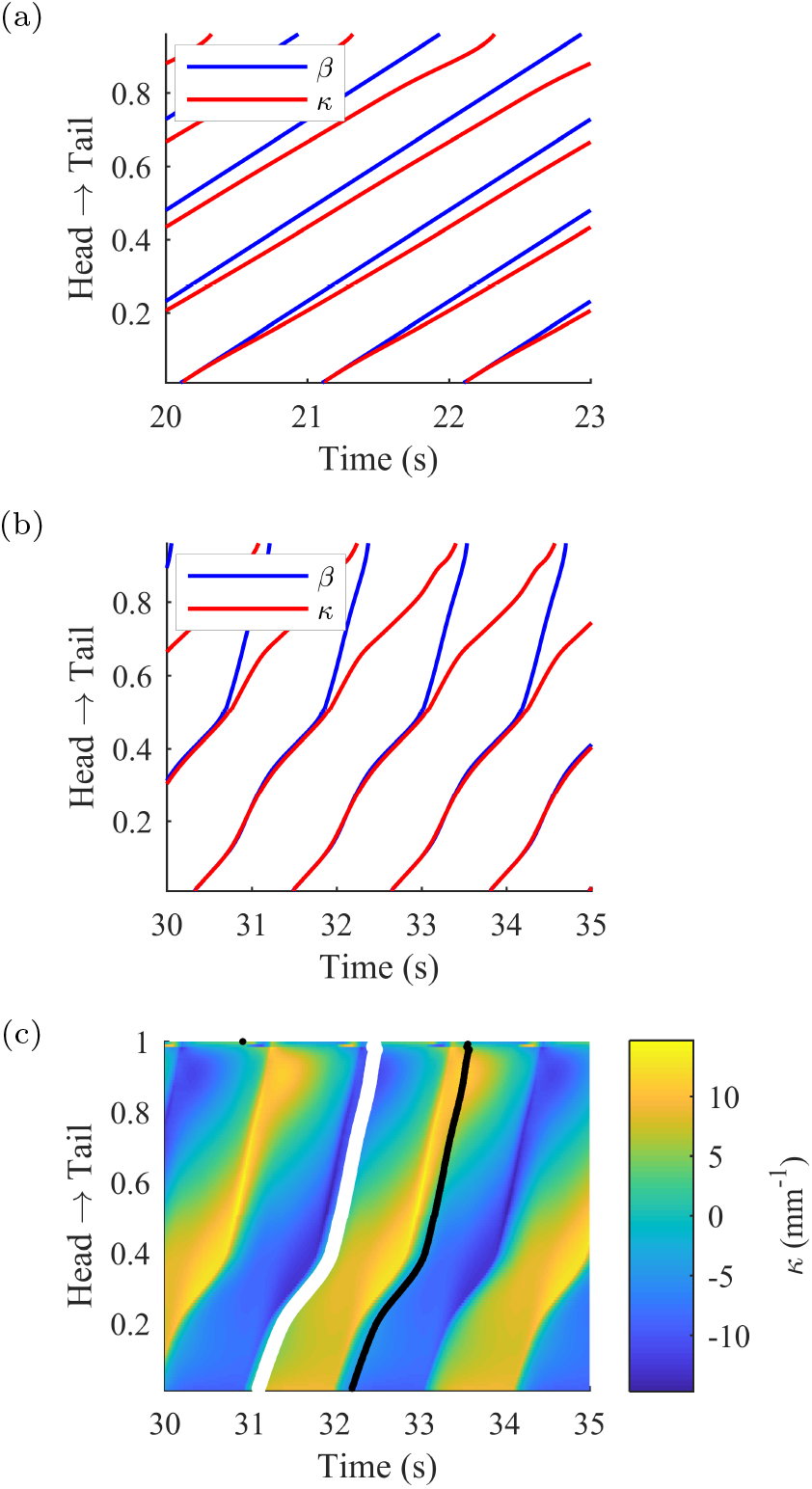
Contours of zero preferred and actual curvatures obtained from kymograms for simulated motion in an agar-like medium and *E* = 100 KPa. (a) Feed forward control model showing linearly growing latency between activation and body bend along the body. (b) Feedback control model: The phase lag remains very small in the anterior half of the body but rapidly increases in the posterior half, lagging behind an accelerated neuromuscular wave speed *β*. (c) Corresponding kymogram of body curvature for the same feedback control simulation. Black and white lines show peak negative and positive levels of muscle activation *β* along the body, respectively.

The corresponding (feedback control) curvature kymogram with superimposed contours of peak (positive and negative) excitation in *β* confirms this observation: the ratio of the body torque to environmental resistance is insufficient for the tail to respond as promptly as the anterior of the worm. In contrast, dorso-ventral switching of the neural activation closely follows the peak curvature of the opposite side all along the body. While our results for the posterior part of the body are not likely to match the curvature dynamics in the worm, they highlight the sensitivity of the system to the exact form of sensory input. In fact, the rich dynamics in the tail demonstrate the tight coupling among kinematic parameters of curvature, wavelength, wave speed and acceleration under feedback control.

Finally, we asked whether a proprioceptively-driven model worm swimming in water will exhibit any neuromechanical phase lags. As expected, we found negligible neuromechanical phase lags in water. These results appear similar both qualitatively and quantitatively to those obtained under feed forward control.

### Locomotion speed is sensitive to both neural control and body properties

As we have seen, both the nature of the pattern generation mechanism and body properties tightly affect neuromechanical phase lags. In lamprey, such phase lags emerge in the tail, when the tail beat frequency is sufficiently high. In that case, one interesting observation is that the emergence of such phase lags in the tail can enhance the efficiency of locomotion.^22^ Another, in general unrelated observation in lamprey and other high Reynolds number swimmers is that there exists an optimal stiffness for generating the highest steady locomotion speed.^22^ *C. elegans* locomotion speed was previously characterized in using the same biomechanical model subject to active forcing^6^ (akin to the feed forward control used here, but lacking a dynamic muscle model). Here, we asked how the nature of the control affects neuromechanical coupling and other locomotion metrics for different values of body stiffness.

We measured the absolute locomotion speed in simulated worms, with Young’s modulus values ranging over eight orders of magnitude, from 10 Pa to 1 GPa. Under feed forward control, the period and frequencies of undulations were manually set (as above), whereas under feedback control, the kinematic parameters arose from the neuromechanical coupling. In our simulation results for agar-like conditions (Fig. 4) there appears to be a threshold in the body stiffness below which the speed rapidly decays. We find that the elasticity threshold val ues are similar for feed forward and feedback control, around 20 KPa. Under feed forward control, however, for *E* ≥ 100 KPa, the speed saturates at under 0.2 mm/sec. This speed dependence on the body stiffness contrasts with our results for feedback control. With propriopceptive entrainment, we find that the elasticity threshold at 20 KPa gives rise to a steep rise in speed which peaks at an unrealistically high speed of about 3 mm/sec for a Young’s modulus of the order of 100 MPa. For Young’s modulus values of 20-100 KPa, the locomotion speeds obtained with feedback control are consistently higher than under feed forward control, but both appear consistent with some observed speeds on agar. Under feedback control, higher, but still realistic speeds of 0.2-0.5 mm/sec on agar fall in the range of 100-200 KPa. The above speeds and optimal body stiffness hold for the choice of kinematic parameters used here. As the waveform of undulations can also affect the speed,^6^ these numbers may differ slightly for variations of the model.

**FIG. 4:**
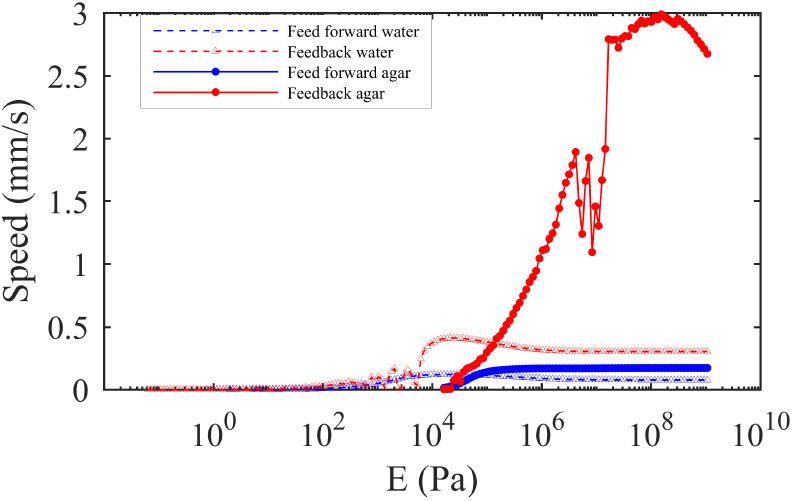
Absolute locomotion speed of simulated worms over a wide range of elasticities. Higher elasticity is required for successful locomotion in agar than in water. Higher speeds are achievable under feedback control than feed forward. Realistic speeds are achieved in the feedback model for a Young’s modulus of *O* ≲100 KPa. Outliers in the plot correspond to simulated trajectories that did not follow a straight line.

Locomotion speeds in water are slightly more difficult to accurately measure experimentally. In our simulations of feed forward control, some (very slow) forward thrust is achieved already with *E* > 100 Pa but the speed peaks at approximately 0.1 mm/sec and slightly decays for high body stiffness values. Simulations with feedback control gave rise to a similar threshold in the Young’s modulus; furthermore, under both feed forward and feedback control, swimming speeds peaked at a similar optimal body stiffness of *E* ≈ 10 KPa. However, once again, we found that feedback driven undulations gave rise to significantly higher locomotion speeds above threshold. In particular, for *E* ≥ 70 KPa speeds of 0.2 mm/sec or higher were observed.

## CONCLUSIONS

In this paper, we set out to compare the mechanical and kinematic manifestations of central pattern generated versus reflex-driven control of undulations. This question has rarely been addressed either in general, or in *C. elegans.* We focused our investigation on two metrics: the neuromechanical phase lag and speed, and found that the range of the worm’s Young’s modulus that gives rise to behaviorally relevant speeds also gives rise to advancing neuromechanical phase lags, at least in the case of feed forward control.

Neuromechanical phase lags are known to exist and affect the locomotion in a variety of undulatory animals, and yet, have not previously been characterized in *C. elegans.* Having checked the entire range of estimated body stiffness in *C. elegans*, we find that unlike fish, the relatively low beat frequencies of the worm combined with the relatively high bulk Young’s modulus preclude the existence of any significant neuromechanical phase lags in water. The biomechanical model used to generate these results lacks any body damping; thus, quantative kinematic parameters of undulations in this regime will likely need revisiting with a more detailed neuromechanical model. As estimates of internal body viscosity in *C. elegans* are relatively low, we expect the key results to hold.

Unlike swimming in water, when crawling on agar, we found that neuromechanical phase lags can arise but depend strongly on the nature of the pattern generation. In a model of feed forward control given by a sine wave that propagates down the animal, we observe potentially significant linearly advancing phase lags along the body of the worm. When the rhythm is entrained by the body curvature, however, as in our model of feedback control, we observe a strong clamping of neuromechanical phase lag, over a wide range of values of body stiffness.

In our model of feedback control, neuromechanical phase lag emerged in the tail, but further work would be required to better understand their origin and physiological significance, if any. In particular, if such phase lags arise in proprioceptively driven locomotion, our model suggests that they likely depend on the form and receptive field of the proprioceptive sensory input. Since mechanical feedback acts to minimize the delay between muscle activation and body bend, different proprioceptive mechanisms or parameters may have the potential to eliminate or facilitate such delays as needed along the body.

## ACKNOWLEDGMENTS

We thank Felix Salfelder for useful discussions and Yang Ding for his thoughts on neuromechanical phases in sandfish. This work was funded by the EPSRC (EP/J004057/1). TR is funded by a Leverhulme Trust Early Career Fellowship.

## APPENDIX: SIMULATIONS AND KINEMATIC PARAMETERS

All simulations were performed for 60 seconds using integration time steps of 0.3 ms. Before any processing, the transients of all simulations were removed manually.

**Phase Lag:** To calculate neuromechanical phase lag *φ*(*u,t*), we took the Hilbert transform of the torque, *β*, and curvature, *κ*, separately using MATLAB’s inbuilt hilbert function. Unwrapping the angles yielded monotonically increasing phases along the body for each point in time, *φβ*(*u,t*) and *φ_κ_*(*u,t*). The neuromechanical phase lag *φ*(*u, t*) was defined as the time averaged phase lag at each body point by *φ_κ_*(*u,t*) – *φβ*(*u,t*).

**Locomotion speed:** To remove side-to-side displacement arising from the undulatory movement, a straight line fit was performed over the trajectory of the midpoint of the worm’s body (u = 0.5) over time. The speed was then defined as the distance traveled along the straight line over the corresponding time interval.

